# Individualized Functional Connectivity-Guided TMS Targeting Theory of Mind Network for Autism Spectrum Disorder

**DOI:** 10.64898/2026.04.09.717580

**Authors:** Na Zhao, Bin Zhang, Xiu-Qin Wang, Hongjian He, Peiying Li, Xian-Wei Che, Robin F.H. Cash, Steven Laureys, Ling Sun, Yu-Feng Zang, Li-Xia Yuan

## Abstract

Transcranial magnetic stimulation (TMS) shows promise in autism spectrum disorder (ASD), but variable outcomes may reflect suboptimal targeting. We developed a functional-connectivity (FC)-guided individualized TMS approach by identifying an ASD-relevant effective region and selecting superficial targets. In a multi-site mega-analysis of Autism Brain Imaging Data Exchange I data (298 ASD, 348 controls), the region with the greatest regional homogeneity (ReHo) abnormality was defined as the effective region. Individualized dorsolateral prefrontal cortex (DLPFC) and inferior parietal lobule (IPL) targets were localized as sites with strongest FC to this region. Group differences, symptom associations, and a six-patient case series were examined. The posterior cingulate cortex (PCC) showed the greatest ReHo abnormality and was implicated in theory-of-mind (ToM) circuitry. PCC-guided targets showed weaker FC in ASD in the right IPL, correlating with Autism Diagnostic Interview social scores; left DLPFC FC differences lacked symptom associations. In the case series, individualized PCC-IPL-guided TMS reduced ToM-related symptoms and Childhood Autism Rating Scale scores. PCC-IPL FC-guided TMS is a biologically informed intervention for modulating ToM circuitry in ASD.

---

Autism spectrum disorder (ASD) is characterized by a constellation of social communication deficits, repetitive and restricted behaviors and interests, and atypical sensory functions manifested in early childhood. It is a pervasive, highly heritable, and heterogeneous neurodevelopmental disorder ^1^. Although many behavioral and pharmacological interventions exist, they are either highly expensive and time-consuming or not effective for the core symptoms ^2,3^. Meta-analyses have revealed that transcranial magnetic stimulation (TMS) is a safe non-invasive neuromodulation technique and demonstrates potential in alleviating ASD-related core deficits ^4,5^. However, the effect size of TMS therapy is moderate and varies widely among studies. This may relate to the imprecise localization of superficial stimulation targets. For example, the stimulation targets in the widely used dorsolateral prefrontal cortex (DLPFC) are typically determined using the “5-cm” method, but this approach misplaces the intended target outside the DLPFC in more than 60% of cases ^6,7^. Individualized precision targeting can ensure appropriate targeting and is a promising approach for enhancing the efficacy of TMS for ASD.

Recent studies have demonstrated the potential to optimize TMS treatment efficacy based on individual-specific functional connectivity (FC) ^8–11^. Individualized precision TMS targeting can be guided with the FC of a core pathological region, also referred as the “effective region” ^8,12^. This region represents a critical abnormal neural node and is consistently implicated in neuropathology ^8,10,11,13^. For many neuropsychiatric disorders, the most implicated (core) pathological region is located in the deep brain ^13–16^. To modulate the activity or connectivity of this deeper structure, interconnected regions at the cortical surface can be identified based on FC and then targeted using TMS. Previous research demonstrates that this approach can modulate the local activity, functional connectivity, effective connectivity, and signal shift of the deep effective region, and may be more effective when this target is individualized according to person specific brain network architecture ^12,17–21^. Recently, personalized amygdala-DLPFC-FC-guided TMS preliminary showed optimized treatment outcomes in young minimally verbal children with ASD ^18^. However, the established core pathological region and optimal corresponding superficial stimulation zone remains unclear. Understanding these aspects is essential for developing individualized TMS therapy for ASD.

In ASD, one candidate effective region is the posterior cingulate cortex (PCC). This is a hub node of the theory of mind network (ToM), which supports mental-state inference and perspective taking-processes often impaired in ASD and closely tied to core social-communication difficulties ^22,23^. The ToM network is strongly spatially overlapping with-but not identical to the default mode network (DMN), sharing core hubs such as the PCC/precuneus, medial prefrontal cortex (mPFC), and angular gyrus/temporoparietal junction ^24,25^. Converging evidence from molecular, structural, and functional imaging studies have revealed that the ASDs exhibit a range of abnormalities in the PCC ^26–29^. Specifically, a significant increase in choline level/GABA_A_ receptor within the PCC was reported ^27,28^, alongside altered cytoarchitecture ^29^ and diminished local activities, such as regional homogeneity (ReHo) and voxel-mirrored homotopic connectivity ^26,30–32^. Collectively, these findings suggested that the PCC may be a core pathological region for ASD. Thus, the PCC is a promising effective region for the treatment of ASD in individualized FC-guided TMS therapy.

Two promising candidate TMS-accessible (i.e. superficial cortical) regions for modulating the PCC, as a downstream effective region, include the DLPFC and the IPL ^4,5,33–35^. The DLPFC is a main node for executive functions, and is demonstrates hyperconnectivity with the ToM in ASD ^36,37^. Stimulating the DLPFC can improve executive function and alleviate comorbidity with depression in ASD ^38^. The IPL is an important component in the ToM, strongly connected with the PCC, and responsible for the theory of mind, social communication, and attention deficits ^35,39^. Previous studies revealed abnormally reduced connectivity of IPL with the PCC, right dorsal premotor cortex, and the cerebellum in ASD ^32,40^; these aspects were strongly correlated with impairments in social-communicative functioning ^40^.

In this study, we aimed firstly to delineate the region within the PCC that is most aberrant in ASD, by performing mega-analysis on a large-scale multi-site dataset. The associated psychological functions of this region were additionally summarized using Neurosynth. Then, individualized potential stimulation targets within the DLPFC and IPL were localized separately according to the strongest FC with the defined deep effective region. Next, group differences in peak FCs within both the DLPFC and IPL were separately estimated between ASD and typically developing controls (TDC). Subsequently, the underlying relationships between these peak FCs and clinical symptoms were evaluated to confirm the optimal superficial functional zone. Finally, we conducted a case series study to empirically validate the feasibility and efficacy of the theoretically optimal individualized FC-guided TMS therapy. Our findings may not only enhance the efficacy of current clinical TMS therapy for ASD, but also offer new insights into individualized TMS therapies for other neuropsychiatry disorders. An overview of the study design and analytic workflow is shown in Fig. 1.

**Fig. 1.**
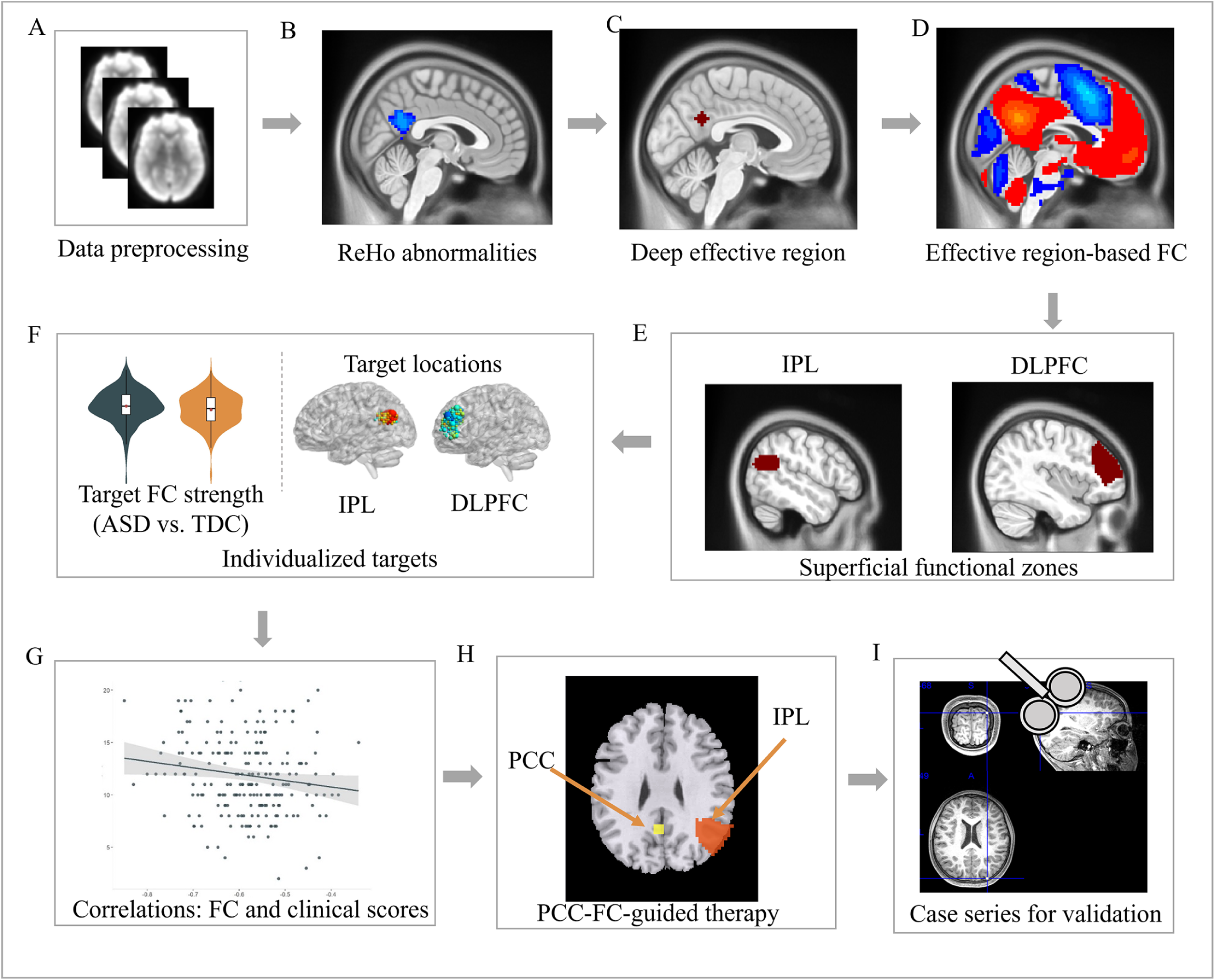
Study design and data analysis overview. A: data preprocessing; B: brain regions with significant ReHo abnormalities in ASD relative to TDCs; C: localizing deep effective region in the PCC based on peak ReHo abnormalities in ASD; D: seed-based FC analysis of the deep effective region; E: segmentation of superficial stimulation cortices including the IPL and DLPFC with watershed algorithm; F: defining individual voxels with peak FC as potential stimulation targets in the bilateral IPL and DLPFC; G: correlation analysis between peak FC strengths and clinical scores. H: the PCC-peak-FC-guided TMS therapy; I: Case series study for validation. ASD: autism spectrum disorder; ReHo: regional homogeneity; TDCs: typically developing controls; PCC: posterior cingulate cortex; FC: functional connectivity; IPL: inferior parietal lobule; DLPFC: dorsolateral prefrontal cortex.

## Results

### Most abnormal ReHo in the PCC

Compared with the TDC group, the ASD group had significantly decreased ReHo in the PCC (*t* = −5.365) and increased ReHo in the left middle frontal gyrus (*t* = 4.190), right superior medial frontal gyrus (*t* = 4.223), and bilateral superior temporal cortices (left: *t* = 4.328; right: *t* = 5.362) (Fig. 2A). Among these regions, the PCC showed the largest difference between ASDs and TDCs. The location (MNI coordinate: [−3, −54, 27]) with the most abnormal ReHo in the PCC was considered to be the core pathological area in ASD and confirmed as the deep effective region for further TMS treatment (Fig. 2B). A meta-analysis based on Neurosynth revealed that the defined effective region was associated with cognitive functions include autographical memory, affective social processing, and mental states (Fig. 2B).

**Fig. 2.**
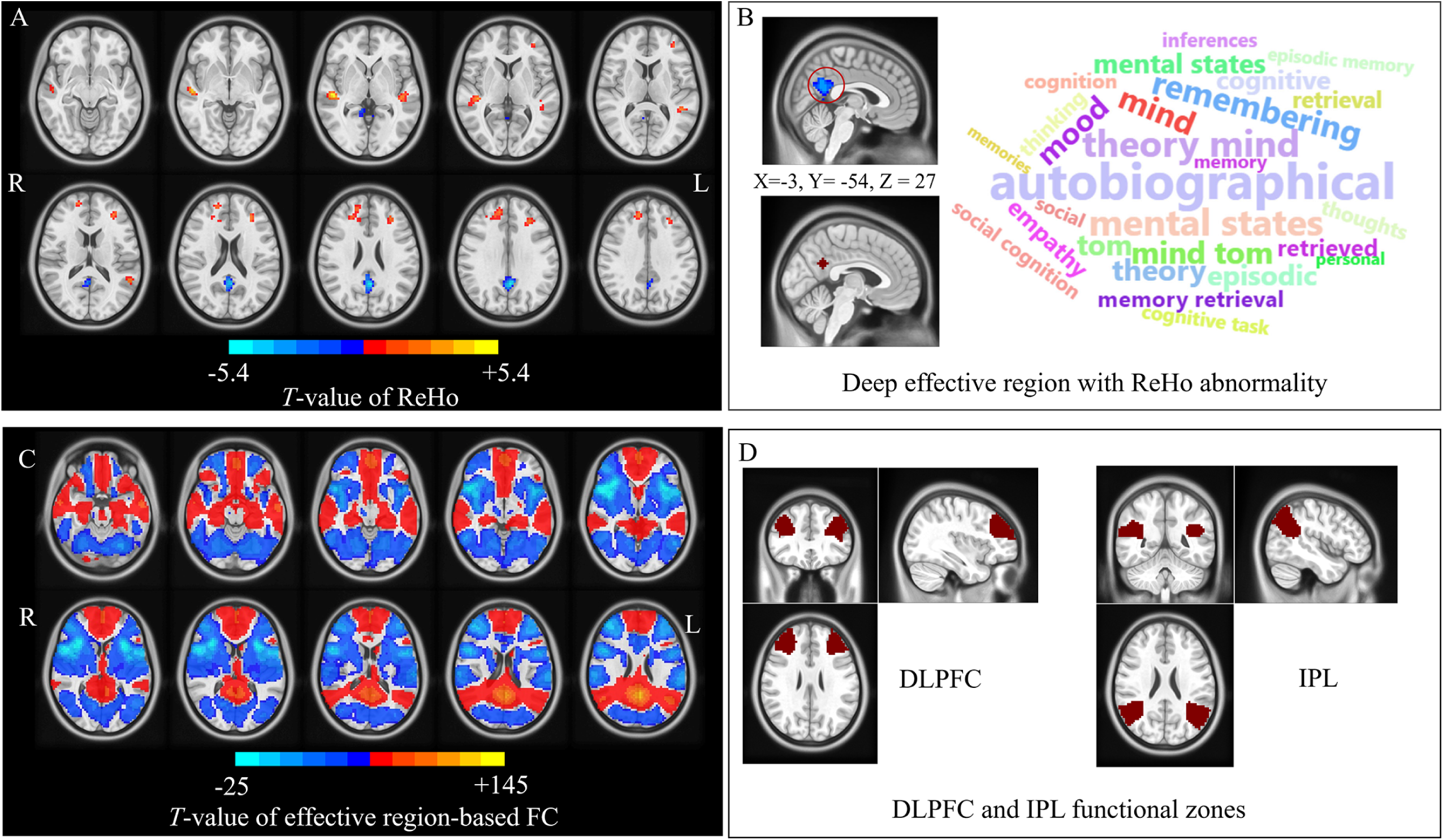
Deep effective region identification and superficial stimulation functional zones. A: Two-sample *t*-test results of ReHo between ASDs and TDCs displayed from −10 mm to +35 mm at axis view; B: the deep effective region in the PCC with peak abnormal ReHo, and the associated psychological functions of the localized deep effective region in the PCC using the Neurosynth database; C: One-sample *t*-test result of the deep effective region-based FC for ASDs displayed from −20 mm to +25 mm at axis view and (D) the potential superficial stimulation zones, i.e., the DLPFC and IPL refined using watershed algorithm. ASD: autism spectrum disorder; TDC: typically developing control; ReHo: regional homogeneity; PCC: posterior cingulate cortex. IPL: inferior parietal lobule; DLPFC: dorsolateral prefrontal cortex; R: the right hemisphere; L: the left hemisphere.

### Functional connectivity of the deep effective region in the PCC

In the seed-based FC analysis in ASD, the deep effective region in the PCC had significant positive connections within the ToM (the bilateral IPL, medial prefrontal cortices, and precuneus) as well as negative correlations with the bilateral temporal, occipital, and middle and lateral prefrontal cortices (Fig. 2C). Of these, the IPL was positively correlated with the defined effective region, whereas most of the DLPFC was negatively correlated with this region (Fig. 2C). The superficial functional zones of DLPFC and IPL refined with watershed algorithm on the group-level seed-based FC were showed in Fig. 2D, which spread over smaller areas relative to that from anatomical atlases (Fig. S1).

### Individualized superficial TMS targets in the DLPFC

The potential superficial targets displayed large variability across individuals and were distributed widely across the bilateral DLPFC (Fig. 3A, Fig. S2). The average scalp depths of individualized superficial targets within the bilateral DLPFC were around 8 mm (Fig. 3B), which indicated that these targets were accessible by TMS. The FC strengths of individualized TMS targets in the left DLPFC (ASD: 0.67 ± 0.15 vs. TDC: 0.70 ± 0.15) were significantly higher in ASDs than in TDCs (Cohen’s *d* = 0.24, *p* = 0.003; 95% confidence interval (CI): [0.08, 0.39]; Fig. 3C). However, there were no significant differences in FC strengths of TMS targets in the right DLPFC. Additionally, no significant correlations were found between FC strengths of TMS targets in the left or right DLPFC and clinical scores.

**Fig. 3.**
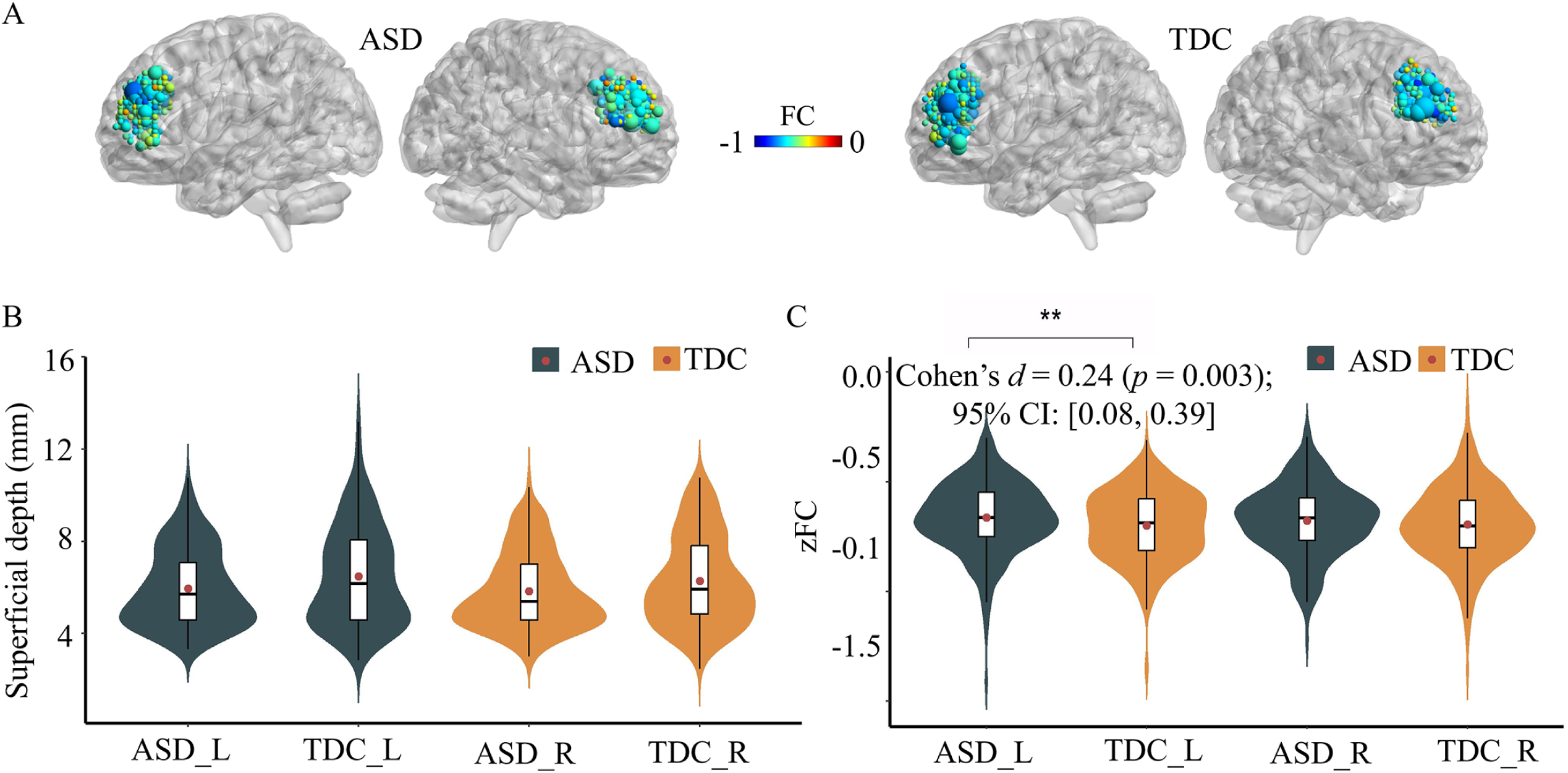
The distribution of individualized potential stimulation targets within the DLPFC (A), their corresponding superficial depth (B), and FC strengths (C). In A, different colors represent the maximum FC strengths and the size of the node indicates the number of duplicate coordinates. ASD: autism spectrum disorder; TDC: typically developing control; DLPFC: the dorsolateral prefrontal cortex.

### Individualized superficial TMS targets in the IPL

Similar to the individualized superficial TMS targets in the bilateral DLPFC, those in the IPL displayed large variability across individuals and were widely distributed (Fig. 4A, Fig. S3). The average scalp depths of individualized superficial stimulation targets within the IPL were around 8-9 mm (Fig. 4B), which were close to those in the DLPFC. The individualized stimulation targets had significantly lower FC strength in ASDs than that in TDCs (ASD: 0.86 ± 0.22 vs. TDC: 0.91 ± 0.22) in the right IPL (Cohen’s *d* = - 0.18, *p* = 0.006; 95%CI: [−0.34, −0.03]; Fig. 4C), while no significant difference in the left IPL. Importantly, the FC strengths of the TMS targets were negatively correlated with ADI social scores in the right IPL (*r* = −0.15, *p* = 0.037; Fig. 4D), while no significant correlations with clinical scores in the left IPL.

**Fig. 4.**
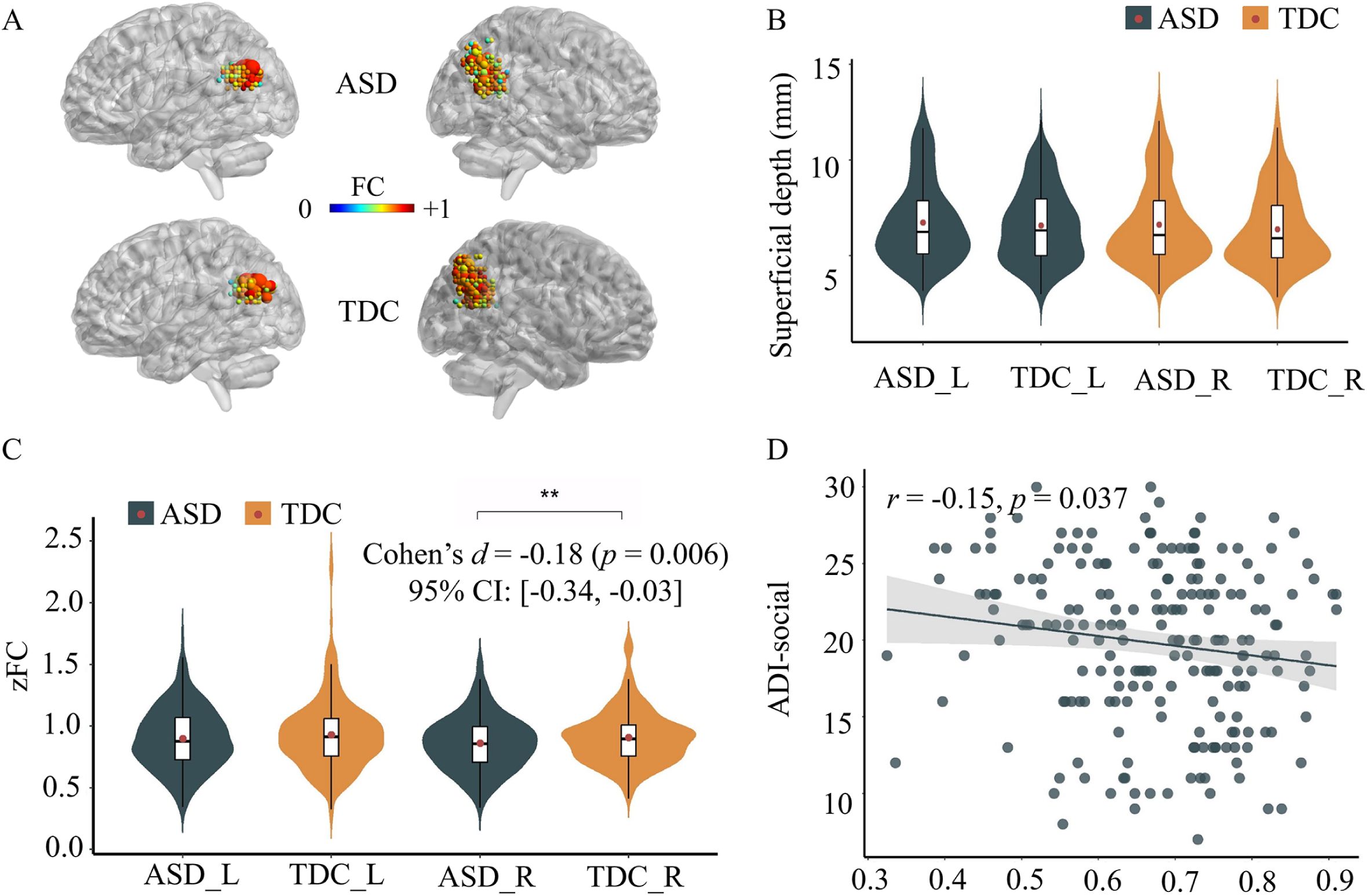
Distribution of individualized potential stimulation targets within the IPL (A), their corresponding superficial depth (B), peak FC strengths (C), and the correlation between peak FC strengths within the right IPL and ADI social scores (D). In A, different color represents the maximum absolute FC strength and the size of the node indicate the number of duplicate coordinates. ASD: autism spectrum disorder; TDC: typically developing control; IPL: inferior parietal lobule; ADI-social: the social sub-scores of the Autism Diagnostic Interview.

### Alleviating social deficits with PCC-IPL-FC guided TMS treatment

After an 8-week TMS therapy, the overall CARS score decreased to 24.8 from 28.2 (*t* = −3.492, *p* = 0.017; Cohen’s *d = -*2.338; 95% CI: [−2.569, −0.224]), presenting a reduction of 12% (Fig. 5, Table S1). Among these patients, two patients showed clinical meaningful improvement after treatment with a 24% and 17% reduction in CARS total score, respectively. Among the subscales, the CARS-3 (Emotional Response) showed the largest improvement with a 28% reduction, and a moderate alleviation was observed in the CARS-8 (Listening Response) and CARS-10 (Fear/ Anxiety), both a 19% decrease (Fig. 5C, Table S1).

**Fig. 5.**
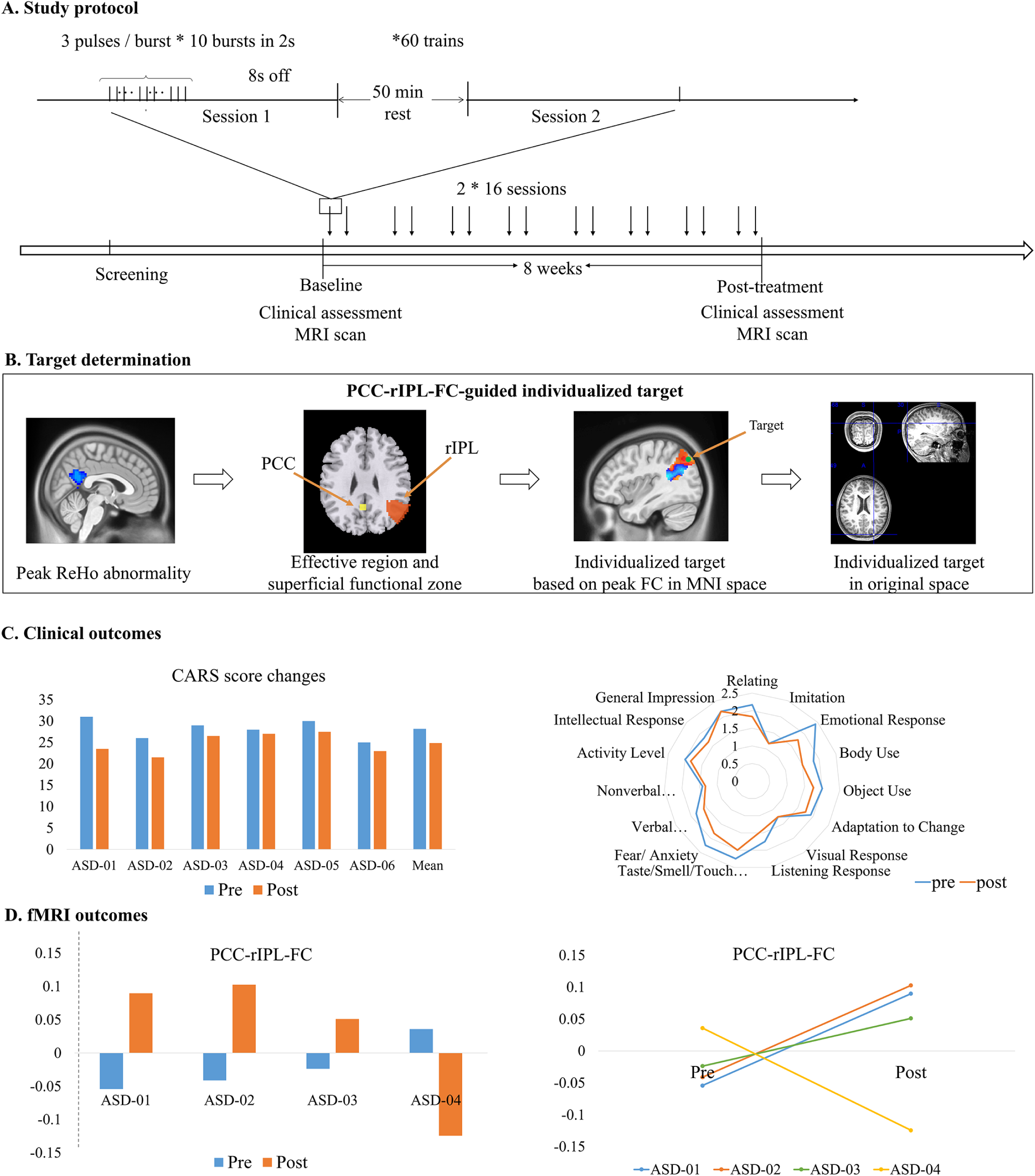
Case series study for validation. A: detailed study protocol; B: PCC-rIPL-FC-guided stimulation target definition; C: CARS reduction after TMS therapy (*t* = - 3.492, *p* = 0.017; Cohen’s *d =-*2.338; 95% CI: [−2.569, −0.224]); D: changes of the PCC-rIPL-FC after TMS therapy. PCC: posterior cingulate cortex; rIPL: right inferior parietal lobule; CARS: Childhood Autism Rating Scale; *: *p*<0.05.

Post-treatment fMRI data were available for four participants. In three individuals (ASD-01, −02, and −03) who showed notable clinical improvement (Table S2), individualized TMS produced a marked increase in FC between the deep effective region and the stimulation functional zone (right IPL) (Fig. 5). In contrast, ASD-04, who exhibited only a minimal 1-point reduction in CARS score, showed a substantial FC decrease from 0.04 to −0.12.

## Discussion

FC-guided individualized TMS has the potent to optimize the treatment efficacy of ASD ^8–11,18^. However, the deep effective region and its corresponding superficial functional zone for the FC-guided therapy for ASD are far from clear. To fill these gaps, we firstly revealed that the PCC had the most atypical ReHo in ASDs than in TDCs and was a core region of neuro-disruption, which is a potential deep effective region for TMS therapy. Then, we demonstrated the superiority of the PCC-IPL-FC guided individualized TMS therapy over the PCC-DLPFC-FC guided intervention by comparing the associations between the personalized targets and clinical symptoms. Finally, a case series study was conducted to further validate the efficacy of the PCC-IPL-FC guided individualized TMS treatment on six ASD patients, and we observed remarkable alleviated social deficits after an 8-week personalized therapy.

### Deep effective region in PCC with the most abnormal ReHo

The PCC is a key node of the ToM and converging evidence from biomedical, structural, and functional studies have revealed abnormalities in PCC in ASD (Libero et al., 2016; Liloia et al., 2023; Oblak, Gibbs, et al., 2011; Oblak, Rosene, et al., 2011). ReHo presented a useful measure to narrow down and constrain the PCC target based on abnormalities in ASD. ReHo measures regional homogeneity in brain activity and has been used to identify functional disruptions in brain disorders, such as Alzheimer’s disease, Parkinson’s disease, major depression disorder, and attention deficit hyperactivity disorder ^17,41–45^. Our results revealed decreased ReHo in the PCC, which is consistent with prior meta-analyses ^26,46,47^. Reduced ReHo has been interpreted to reflect higher functional segregation and reduced complex information processing ^48,49^, in line with hallmark features of cognitive inflexibility and impaired social-emotional processing in ASD. Furthermore, aligning with previous research, Neurosynth-based meta-analysis indicated psychological functions of the deep effective region include autographical memory, theory of mind, affective social processing, and mental states ^50–53^, which were impaired in ASD. Thus, the region with the most abnormal ReHo in PCC is a promising effective region for the treatment of ASD in individualized FC-guided TMS therapy.

### PCC-FC-guided individualized TMS treatment for ASD

Individualized FC-guided TMS therapy has been illustrated to be an effective new method for treating various neuropsychological disorders, such as depression, ASD, and Tourette’s syndrome ^8,17,18,21,54,55^. In ASD, a personalized amygdala-DLPFC-FC-guided TMS therapy preliminary demonstrated its superior clinical efficacy in young minimally verbal children with ASD ^18^ compared to targeting determined with EEG F3 site. Our results revealed that the PCC-FC-guided individualized TMS sites scattered throughout the whole DLPFC and IPL, and this large variability among participants echoes the essentiality for individualized targeting.

Moreover, substantial evidence from previous researches and our results have highlighted impaired local activity within the PCC and its functional connectivity with the frontal, parietal and temporal cortices, particularly the ToM areas including the medial prefrontal cortex and IPL ^31,40,56–58^. Additionally, the abnormal connectivity of the PCC was closely associated with deficits in face memory, emotion perception, praxis and social-communicative skills ^26,40,59,60^. As TMS holds promise for modulating the brain activity of the deep effective region and its related network ^8,9,12,17–21,61^, the PCC-FC-guided individualized TMS might effectively alleviate ASD-related symptoms, such as social functioning and emotional processing.

### Individualized TMS targets in the IPL

According to our previous review on non-FC-guided TMS treatment for ASD, the DLPFC and IPL are the most commonly used superficial targeting functional zones ^33^. Our results showed that the FC strengths of individualized TMS targets demonstrated group differences in both the right IPL and the left DLPFC, while only the connectivity between the PCC and right IPL was negatively correlated with the ADI social score. These findings indicated that the FC between the PCC and its optimized superficial targets in the right IPL is a potential ASD biomarker, which aligns with prior result that weaker connectivity between the PCC and right IPL was closely associated with poorer praxis and social-communicative skills ^40^. Furthermore, previous literatures demonstrated that TMS can modulate not only the local activity of the deep effective region, but also its FC strengths with superficial targets per se ^9,12,17,18,21^. Thus, it is reasonable to speculate that PCC-FC-guided stimulation can modulate these abnormal peak FCs and alleviating social deficits. Additionally, the PCC has stronger FC with the IPL than the DLPFC. Similar to the PCC (the potential effective region in the current study), the IPL is part of the ToM ^62^ and is involved in theory of mind and social communications ^35,39^. While, DLPFC belongs to the executive control network and is important for executive function ^38^. Taken together, with the PCC as the deep effective region, the right IPL is theoretically a better superficial stimulation zone than the DLPFC, and PCC-IPL-FC-guided individualized TMS may help relieve social impairment in ASD.

### Alleviated social deficits after PCC-IPL-FC guided TMS treatment

In the case series performing the PCC-IPL-FC-guided TMS therapy, all six ASD patients demonstrated a reduction in CARS score, including sub-symptoms in listening response, emotional response, and fear or anxiety. Pre-to-post changes in symptom severity were significant across the group. These findings are promising and provide preliminary evidence for success of this strategy. Further comparator-controlled research is warranted to determine the relative efficacy of this approach compared to more conventional targeting methods. These findings align with previous non-FC-guided TMS case studies ^34,35^. For instance, Yang et al. reported alleviation of social-related symptom in a cohort of 11 ASD patients following TMS therapy on the IPL ^35^. Moreover, our neuroimaging analyses showed that the therapeutic effects were tied to the normalization of aberrant FC. After the 8-week PCC-IPL-FC-guided TMS treatment, four of the six patients underwent follow-up scans. Three out of four (75%) patients exhibited both increased PCC-right IPL connectivity and remarkable clinical improvement. In contrast, the remaining patient who showed only a minor CARS reduction demonstrated a decrease in FC. These findings would be consistent with the relation between FC restoration and clinical response.

These results provide empirical evidence that PCC-IPL-FC-guided individualized intervention alleviates ASD social-related symptoms through specifically targeting and enhancing the pathologically reduced FC in affected regions. This mechanism is consistent with previous studies that FC-guided TMS treatment could modulate the FC of the deep effective region ^9,12,18,63^. Collectively, the case series performing PCC-IPL-FC-guided individualized TMS intervention preliminarily and empirically showed its feasibility and potential efficacy in social symptom alleviation.

### Limitations

Our study also exists some limitations. First, 87.9% (i.e. n of N) of the included ASD sample were males, while sex discrepancies are crucial for brain development during ASD progression. Thus, future studies focusing on females are necessary for individualized TMS therapy. Second, the mega-analysis of a large, multi-center dataset with higher quality control would allow for greater generalization of the results. In the present analysis, we excluded many samples because of various heterogeneities in the ABIDE I data. Third, the superiority of the PCC-IPL-FC-guided therapy for ASD-related social symptoms was only validate on open-label case series, its clinical efficacy needs more validation in future empirical studies.

## Conclusions

In this study, we developed an innovative individualized PCC-IPL-FC guided TMS therapy based on a large multi-site dataset targeting the ToM, which was further validated with a case series showing notable clinical alleviation on ASD-related emotional and social processing symptoms and normalization of the pathologically reduced PCC-IPL FC. This work paves the way for biologically informed and precision therapeutic strategies for individuals with ASD.

## Methods

### Participants

We used the data from Autism Brain Imaging Data Exchange I (ABIDE I) (http://fcon_1000.projects.nitrc.org/indi/abide/), comprising 1112 participants (539 ASDs and 573 TDCs) from 17 sites. Participants were excluded according to the criteria detailed in the Supplementary materials (Fig. S4) and our previous researches ^64,65^. Finally, 646 participants comprising 298 ASDs and 348 TDCs from 14 sites were analyzed (Table S3). All data collected from each site were approved by the local Institutional Review Boards and written informed consent were obtained for all participants (http://fcon_1000.projects.nitrc.org/indi/abide/). The demographic information differences between the ASD and TDC were assessed with two-sample *t*-test or chi-square test as appropriate. These comparisons were two-tailed and the significance threshold was set as *p* < 0.05.

### Data processing

The fMRI data preprocessing was conducted using DPABI 5.1 ^66^ (http://rfmri.org/dpabi) and SPM12 (https://www.fil.ion.ucl.ac.uk/spm/software/spm12/) according to our previous researches ^64,65^ as follows: 1) removing the first 10 volumes; 2) correcting slice timing; 3) realigning for head motion correction; 4) spatial normalizing to Montreal Neurological Institute (MNI) space; 5) removing linear detrend; 6) regressing out head motion parameters based on Friston-24, white matter signal, cerebrospinal fluid signal, and global mean time series; and 7) band-pass filtering (0.01**-**0.08 Hz). Subsequent analyses were then performed as described in Fig.1.

### Detection of the core pathological region in PCC using ReHo

ReHo is a voxel-wise metric that describes the brain activity synchronization of a given voxel with its neighboring voxels ^67^, which is widely used to localize the detailed core pathological region in neuropsychiatry disorders ^41,42^. After preprocessing, ReHo was calculated and spatially smoothed using a Gaussian kernel with full width at half maximum of 6 mm. For standardization, the mean ReHo was obtained by dividing its whole brain mean value. Then, ComBat (https://github.com/Jfortin1/ComBatHarmonization) was used to remove site-related variance produced by differences in collection time, imaging parameter, machine performance, and participant character across multi-sites ^68,69^.

The group differences of ReHo between ASD and TDC were analyzed using independent two-sample *t*-tests (Gaussian random field [GRF] correction, cluster *p* < 0.05; voxel *p* < 0.001) with the mean framewise displacement (mFD) and full-scale intelligence quotient (FIQ) scores as covariates. Then, the brain area with the most abnormal ReHo activity was defined as the core pathological region and the deep effective region for TMS. Furthermore, the associated psychological functions of the deep effective region were summarized using Neurosynth (https://neurosynth.org/locations/).

### Whole brain functional connectivity of the deep effective region

Seed-based FC was applied to obtain the whole brain functional connectivity of the deep effective region. Firstly, the seed was defined as a 6-mm sphere centered at the deep effective region and the mean time course was computed by averaging the time courses of all voxels in the seed. Then, Pearson’s correlation coefficients between the mean time course and those of other voxels across the whole brain were calculated as the FC values. Finally, the FC values were Fisher’s *Z*-transformed to increase the normalization of the distribution, and one-sample *t*-test was performed to detect the superficial cortices with group-level significant FCs (GRF correction, cluster *p* < 0.05; voxel *p* < 0.001)

### Definition of FC-guided individualized superficial TMS targets

Firstly, the DLPFC and IPL zones were initially defined according to previous studies ^70–72^. Specifically, the left DLPFC was defined as the interaction area of 20 mm radius spherical ROIs centered at the left Brodmann area (BA) 9 (MNI: −36, 39, 43), BA 46 (MNI: −44, 40, 29), the “5-cm” TMS site (MNI: −41, 16, 54), and F3 Beam group average stimulation target (MNI: −37, 26, 49) ^70,72^. The right DLPFC was obtained by mirroring the left DLPFC. The IPL was defined as the overlap of the IPL regions within the Oxford-Harvard cortical grey matter atlases and the angular cortex within the Automated Anatomical Labeling template ^71^. Then, the DLPFC and IPL zones were further refined by restricting only one peak in the cluster with the watershed algorithm on the group-level significant FC maps of the defined effective region ^73^.

The personalized superficial stimulation targets were defined as locations owing strongest FC strengths within the left DLPFC, right DLPFC, left IPL, and right IPL, separately, for each individual. Then, the superficial depths of these individualized stimulation targets were calculated, which is essential for assessing their TMS accessibility as the commonly used figure-eight coil can only focus on regions at a depth of 2-3 cm ^74^.

### Associations between individualized TMS targets and clinical scales

Firstly, to detect the group differences in the connectivity strength and locations of the individualized stimulation targets between ASD and TDC, general linear regression analyses were conducted with mFD and FIQ as covariates. Then, to explore the relationship between individualized stimulation targets and clinical symptoms, we calculated Pearson’s correlation coefficients between peak FC strengths in both the bilateral DLPFC and IPL and clinical scores, including those of the ADI, Autism Diagnostic Observation Schedule, Social Responsiveness Scale, and Social Communication Questionnaire. The significance threshold was set as *p* < 0.05.

### Case series study for validating the individualized FC-guided TMS therapy

As our result revealed that the FC strengths of the individualized TMS targets in the IPL in ASD significantly distinguished from that in TDC and negatively correlated with the clinical social score, while no significant correlation was found between the individualized TMS targets in the DLPFC and clinical scores. Therefore, we conducted a case study to validate the feasibility of the individualized TMS therapy based on PCC-IPL-FC-guided stimulation target. Six participants (11.1 ± 4.5 years old, 5 males) diagnosed as ASD was recruited. A written consent was obtained from their guardians prior to the study. This case series study was approved by the Ethic Committee of Tianjin Anding Hospital (No.: 2024-56).

These participants performed individualized intermittent theta burst stimulation (iTBS) therapy based on the PCC-IPL-FC-guided target. Detailly, the resting motor threshold was estimated before the iTBS stimulation. Subsequently, the individualized iTBS therapy was delivered using a MagPro X100 stimulator with MagOption (MagVenture, Denmark) and a Figure-8-coil (B65 Coil), facilitated by the Localite TMS navigator. Specifically, the iTBS parameters were as below: 50-Hz bursts of three pulses (100% the resting motor threshold) with bursts repeated at a frequency of 5 Hz, bursts presented in 10-sec cycles consisting of 2-sec stimulation and 8-sec rest, 60 cycles session run consisting of 1800 pulses. A total of 32 sessions were performed with 2 sessions carried out a day with 50-minute interval and twice a week on the weekend.

To define individualized TMS target, a resting state fMRI and a high-resolution 3D T1 images were acquired on a 3.0 T Prisma scanner. Resting state fMRI were scanned with following parameters: repetition time = 800 ms, echo time = 30 ms, slice thickness = 2 mm, a total of 72 slices, matrix size = 104 × 104, field of view = 208 × 208 mm^2^, flip angle = 56°, frame = 450. The resolution of 3D T1 imaging was isotropic 0.9 mm. The fMRI data was firstly preprocessed in the same way as described in section 2.2. Then, a spherical seed centered at the deep effective region (MNI coordinates: [−3 −54 27]) with a radius of 6 mm was defined. Subsequently, the seed-based FC was performed in the predefined IPL mask in section 2.5. The therapeutic effect of the PCC-IPL-FC-guided TMS therapy was evaluated using Childhood Autism Rating Scale (CARS) by a professional psychiatrist before and after treatment. Furthermore, we evaluated the alterations of the FC between the deep effective region and the right IPL after TMS therapy.

## Data availability

ABIDE I data are publicly available from the Autism Brain Imaging Data Exchange repository. De-identified data generated during the case series are not publicly available due to participant privacy and ethics restrictions, but may be available from the corresponding authors on reasonable request and with approval from the relevant ethics committee.

## Acknowledgments

We thank Bronwen Gardner, PhD, from Liwen Bianji (Edanz) (www.liwenbianji.cn/), for editing the English text of a draft of this manuscript. We also thank Qiuping Ding from the Center for Brain Imaging Science and Technology, Zhejiang University, for her technical assistance with the MRI scanning.

This work is supported by National Natural Science Foundation of China (No. 82301742), the Medical and Health Science Program of Zhejiang Province (No. 2025HY0678, 2025HY0686), Start-up Research Fund of Hangzhou Normal University (HYQD-2025-457), the Construction Fund of Key Medical Disciplines of HangZhou (No. 2025HZGF02), the Australian National Health and Medical Research Council (Emerging Leadership Investigator grant No. 2017527), the Tianjin Education Commission Research Project (No. 2024ZD041), the Tianjin Health Research Project (Grant No. TJWJ2023QN086), and Collaborative Innovation Center of Hebei Province for Mechanism, Diagnosis and Treatment of Neuropsychiatric Diseases.

## Author contributions

**Na Zhao:** Conceptualization, Methodology, Visualization, Writing-original draft, Funding acquisition**; Bin Zhang:** Data collection, Formal analysis, Validation; **Xiu-Qing Wang:** Formal analysis, Validation; **Hong-Jian He:** Writing-Reviewing and Editing; **Peiying Li:** Data collection, Formal analysis; **Xian-Wei Che:** Writing-Reviewing and Editing; **Robin F.H. Cash:** Writing-Reviewing and Editing; **Steven Laureys:** Writing-Reviewing and Editing; **Ling Sun:** Conceptualization, Data collection, Visualization; **Yu-Feng Zang:** Conceptualization, Supervision, Funding acquisition, Writing-Reviewing and Editing; **Li-Xia Yuan:** Conceptualization, Methodology, Visualization, Funding acquisition, Writing-Reviewing and Editing.

## Competing Interest

None.

## Notes

### Competing Interest Statement

The authors have declared no competing interest.

